# Concurrent maternal stress and THC exposure during pregnancy alters adolescent behavioral outcomes and corticolimbic molecular programs

**DOI:** 10.1101/2025.06.26.661775

**Authors:** Jimmy Olusakin, Mahima Dewan, Atul Kashyap, Daniela Franco, Gautam Kumar, Miguel A. Lujan, Katrina S. Mark, Joseph Cheer, Mary Kay Lobo

## Abstract

Cannabis use during pregnancy is increasing, often to alleviate stress and anxiety, yet the long-term effect of prenatal cannabis exposure alone or in combination with psychosocial stress on offspring neurodevelopment or maternal behaviors remains unclear. Here, we developed a translational rodent model combining prenatal Δ⁹-tetrahydrocannabinol (THC) exposure with chronic psychosocial stress using the maternal witness defeat stress (MWDS) paradigm. Pregnant C57BL/6 mice were exposed to MWDS from gestational day (GD) 3-12 and received daily subcutaneous THC (2 mg/kg) or vehicle until birth. All exposure groups showed impaired maternal behavior, with negative postnatal outcomes and caregiving, with additive effects observed in the combined exposure group. In adolescence, male and female offspring exhibited exposure-specific behavioral alterations. Prenatal stress and combined exposures led to increased anxiety-like behavior and reduced motivated behavior in both sexes, while THC alone primarily impacted female self-care and social behavior. Transcriptomic profiling of the prefrontal cortex (PFC) and nucleus accumbens (NAc) of adolescent offspring revealed sex- and region-specific gene expression changes across all exposure groups. Prenatal THC-, stress-, and combined exposures each altered distinct molecular pathways related to mitochondrial function, synaptic organization, and glial signaling. Comparative analysis with a perinatal fentanyl model revealed shared transcriptional substrates involved in synaptic signaling and circadian regulation. These findings indicate that THC and stress independently and additively impair maternal behaviors with lasting neurodevelopment signatures in offspring.

## INTRODUCTION

The increasing legalization of cannabis across the United States has led to a surge in its use among vulnerable populations, including pregnant people, many of whom cite its anxiolytic and stress-relieving effects as their reasons for continued use during pregnancy (1,2). While cannabis may confer short-term anxiolytic effects, its long-term consequences on maternal health, maternal-infant bonding, and offspring neurodevelopment are not well understood. The placental barrier serves as a critical checkpoint, protecting the fetus from maternal glucocorticoids that may ensue from acute stressors (3,4). Chronic maternal stress can compromise placental protection, increasing fetal exposure and altering inflammatory responses in the placenta and fetal brain (5–7). Additionally, prenatal stress (PNS) exposures can disrupt the fetus’s endocannabinoid (eCB) system, crucial in shaping synaptic plasticity, stress regulation, and emotional behaviors (8–10). As both PNS and prenatal cannabis exposure (PCE) independently alter brain development, their combination may cause cumulative or synergistic neurodevelopmental disruptions in fetal brain development, particularly in brain regions involved in emotional regulation, motivation, and reward processing.

The prefrontal cortex (PFC) and nucleus accumbens (NAc) are central to cognition, emotional regulation, and reward-driven behaviors, and are both modulated by the eCB system (11,12). Studies have demonstrated that early-life stress and cannabinoid exposure result in measurable neuroanatomical changes, including reduced cortical thickness, synaptic loss, and dendritic atrophy in the PFC, alongside decreased dopamine receptor expression and sensitivity in the NAc (13–16). While such alterations may not manifest as overt behavioral symptoms immediately after birth, they may predispose offspring to psychiatric disorders later in life (17–19). Therefore, there is a dire need to identify molecular substrates that could provide insights into mechanisms of endocannabinoid actions to inform novel therapeutics.

In the present study, we investigated whether additive exposure to Δ9 tetrahydrocannabinol (THC), the main psychoactive component of cannabis, modulates maternal behavioral responses to psychosocial stress and whether these combined exposures disrupt offspring development. To model maternal stress, we employed the chronic witness defeat stress (CWDS) paradigm. CWDS is a translationally relevant paradigm used to induce psychological stress (20,21) in rodents without direct physical harm. This model mimics vicarious trauma relevant to aspects of chronic stress in pregnant people (22,23). Using this approach, pregnant dams were subjected to a 10-day CWDS from gestation onset, which we term maternal witness defeat stress (MWDS), while receiving moderate daily doses of THC via subcutaneous injections (24). This experimental design allowed us to model the co-occurrence of prenatal stress and cannabis exposure in a controlled manner and assess their cumulative effects on maternal behavior pre-birth, maternal care after birth, and offspring behaviors and neurobiological outcomes. Immediately following cessation of MWDS, dams were assessed for anxiety-like behavior to evaluate the emotional impact of combined prenatal stress and THC exposure. To uncover the molecular substrates underlying these phenotypes, we performed transcriptomic profiling of the PFC and NAc. Finally, we compared these data to published transcriptomic analysis of perinatal fentanyl exposed (PFE) mice, shown to exhibit sex-specific affective disturbances (25). Our study provides mechanistic insight into how prenatal environments program long-term vulnerability to emotional and motivational dysfunction.

## METHODS AND MATERIALS

### Experimental Subjects

All experiments were performed in accordance with the Institutional Animal Care and Use Committee Guidelines at the University of Maryland School of Medicine (UMSOM) and in accordance with NIH guidelines for the use of laboratory animals. Mice were given food and water ad libitum and housed in the UMSOM vivarium on a 12-h light/dark cycle. Experimental mice were 8-9 weeks old male chronic social defeat stress (CSDS) and pregnant females (MWDS) C57BL/6 mice. CD-1 retired breeder males (>4 months) were used as the aggressors for CSDS/MWDS. Mice were purchased from Charles River Laboratories and acclimated in the vivarium for at least one week. Dams were randomly assigned to four groups across three cohorts (6–8 dams each), with data combined for analysis (Fig. 1B-I). The groups included: (I) dams receiving vehicle injections without stress (vehicle-exposed; n = 20), (II) dams receiving THC injections without stress (THC-exposed; n = 21), (III) dams receiving vehicle injections following MWDS (stress-exposed; n = 22), and (IV) dams receiving THC injections following MWDS (THC/stress-exposed n = 22). Adolescent behaviors included mice from no more than 3 pups/litters/sex/condition.

**Figure 1.**
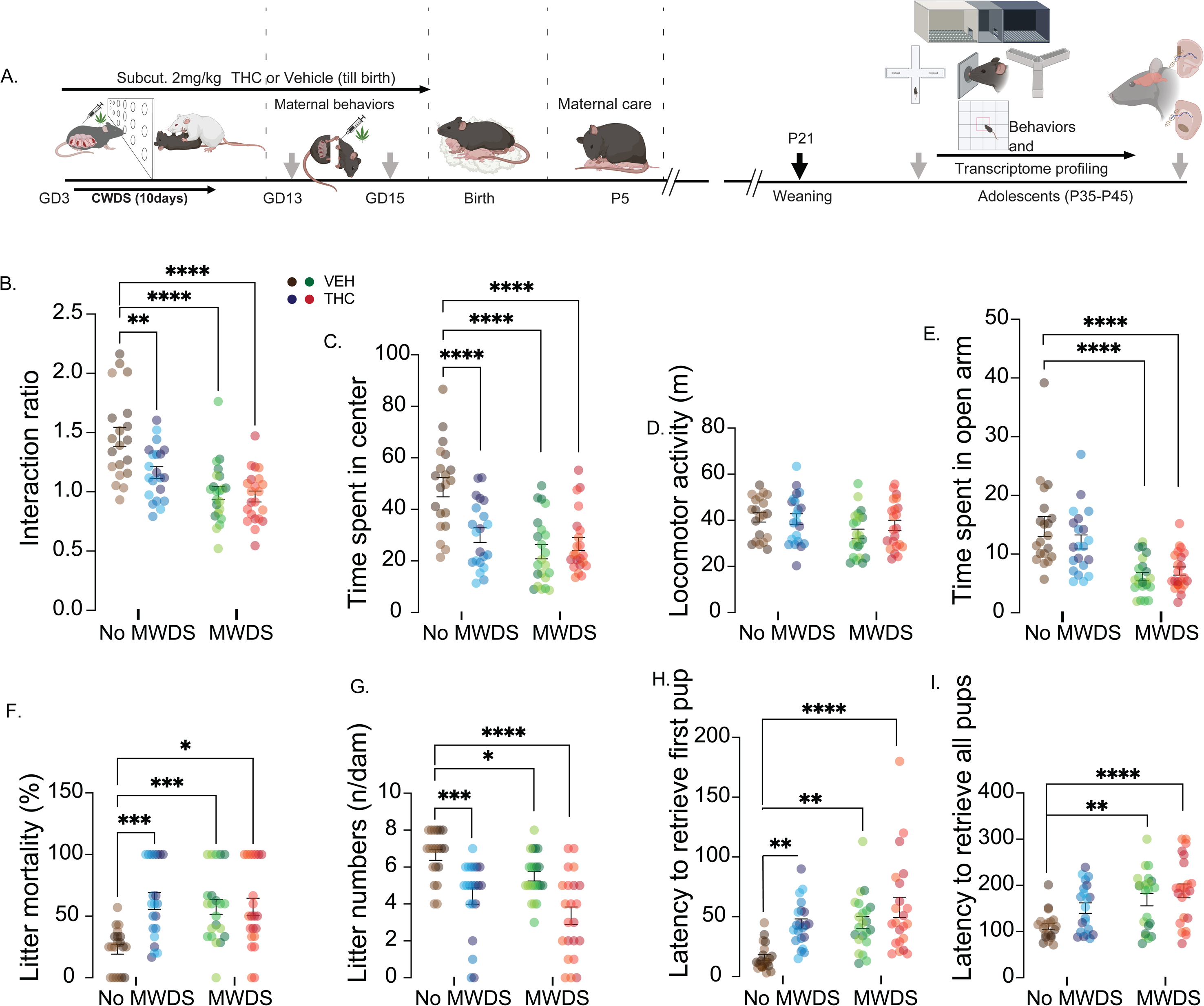
Additive THC exposures to maternal stress heightens negative outcomes and maternal care behaviors. (A) Timeline experimental procedures. Pregnant C57BL/6 dams underwent 10days of MWDS from GD3 followed by concurrent subcutaneous injections of 2 mg/kg, 2 ml/kg THC. Following cessation of MWDS, dams underwent SI, OFT and EPM behaviors. After birth and cessation of THC administration, on postnatal day 5, pup retrieval test was conducted. At postnatal day 35, adolescent mice underwent sequential behavioral paradigms – SI, OFT, EPM, splash test and Ymaze barrier testing. (B-I) Maternal behavioral tests, included 4 groups – control group (brown), THC-exposed group (blue), stress-exposed group (green), THC/stress-exposed group (red). Behavioral testing was conducted in 3 independent cohorts with 6-7 dams/cohort, denoted by variations in color groups. In the SI test (Fig. 1B), two-way ANOVA revealed a significant MWDS × THC interaction (F_1,81_ = 5.184, p = 0.0254), with main effects of MWDS (F_1,81_ = 32.86, p < 0.0001) and THC (F_1,81_ = 7.917, p = 0.0061). Post hoc tests indicated reduced interaction times in THC-exposed (p = 0.0019), MWDS-exposed (p < 0.0001), and THC/MWDS-exposed dams (p < 0.0001) relative to vehicle-treated controls. In the OFT (Fig. 1C), there was also a significant MWDS × THC interaction (F_1,81_ = 13.23, p = 0.0005), and main effects of MWDS (F_1,81_ = 23.12, p < 0.0001) and THC (F_1,81_ = 6.906, p = 0.0103). All exposure groups spent less time in the center zone, consistent with increased anxiety. Locomotor activity (Fig. 1D) showed no significant interaction (F_1,81_ = 1.113, p = 0.2), but a main effect of MWDS was detected (F_1,81_ = 5.073, p = 0.02), with a trend toward reduced distance traveled (p = 0.058). Total locomotion was not changed during the SI test (Figure S1B), suggesting reduced center time in OFT was not attributable to global hypoactivity. On day 3, dams were tested on the EPM (Fig. 1E), and two-way ANOVA showed no interaction (F_1,81_ = 2.622, p = 0.1093), but a robust main effect of MWDS (F_1,81_ = 37.45, p < 0.0001). Post hoc analysis showed MWDS- and THC/MWDS-exposed dams both spent significantly less time in open arms (p < 0.0001) compared to vehicle group, while THC-exposed dams did not differ from vehicle group (p = 0.2), further highlighting stress as the primary driver of anxiety-related phenotypes during pregnancy. Postnatal assessments on day 5, revealed a significant MWDS × THC interaction on litter mortality (F_1,81_ = 13.43, p = 0.004), a main effect of MWDS (F_1,81_ = 4.792, p = 0.0315), but no main effect of THC (F_1,81_ = 3.863, p = 0.0528) (Fig. 1F). Post hoc analysis showed increased mortality in all exposed groups: THC (p = 0.0006), MWDS (p = 0.0003), and THC/MWDS (p = 0.0136). Litter size (Fig. 1G) showed no interaction (F_1,81_ = 0.0124, p = 0.9115), but main effects of MWDS (F_1,81_ = 8.422, p = 0.0048) and THC (F_1,81_ = 32.6, p < 0.0001). Post hoc comparisons showed reductions in THC-exposed (p = 0.0001), MWDS-exposed (p = 0.0373), and THC/MWDS-exposed dams (p < 0.0001) relative to controls. Pup retrieval test (Fig. 1H–I) was done immediately following postnatal assessment and revealed no MWDS × THC interaction in latency to retrieve the first pup (F_1,81_ = 1.852, p = 0.1774) or all pups (F_1,81_ = 0.7164, p = 0.3998), but main effects of MWDS (first pup: F_1,81_ = 14.68, p = 0.0003; all pups: F_1,81_ = 15.22, p = 0.0002) and THC (first pup: F_1,81_ = 13.17, p = 0.0005; all pups: F_1,81_ = 5.739, p = 0.0189) were detected. THC-, MWDS-, and THC/MWDS-exposed dams exhibited significantly prolonged latencies in retrieving the first (THC: p = 0.0025; MWDS: p = 0.0014; THC/MWDS: p < 0.0001) and all pups (MWDS: p = 0.0038; THC/MWDS: p < 0.0001) compared to controls. No significant latencies in retrieving all pups in the THC-exposed dams (p = 0.07). Vehicle-exposed group: n = 20; MWDS group: n = 22; THC-exposed group: n = 21; THC/MWDS: n = 22. MWDS, maternal witness defeat stress; *\*\*p* = 0.01, *\*\*\*p* = 0.001, *\*\*\*\*p* < 0.0001.

### Behavioral Testing

#### Maternal witness defeat stress

The chronic witness defeat stress (CWDS) paradigm (20) was adapted for use in pregnant dams and referred to herein as MWDS. Pregnant dams were housed on one side of a perforated divided cage to witness a 10-minute resident-intruder interaction. The CSDS mouse was then housed opposite the resident, and the MWDS dam was removed and housed opposite to a novel CD-1 aggressor. After 24 h of sensory interaction, the CSDS mouse was defeated by a new CD-1 resident while a different MWDS dam witnessed the agonistic interaction. This process was repeated daily for 10 days with new CD-1–CSDS–MWDS pairings to prevent habituation. Unstressed vehicle and THC dams were pair-housed across perforated dividers in cages containing woodchip bedding for 10days.

#### Three-chamber social interaction

The apparatus consisted of a 60 x 40 x 22 cm arena with white walls and floor, divided into 3 equal chambers (20 x 40 cm) by clear acrylic walls. Wire mesh cups were placed in each outer chamber, while the central chamber remained empty. During the habituation phase, the mouse was placed in the central chamber and allowed to explore the arena containing two empty cups for 5 minutes. Subsequently, a novel, sex-matched conspecific was placed under one of the mesh cups, and the mouse was allowed to explore for an additional 5 minutes (21). Time spent in each outer chamber was recorded using ANY-maze tracking software (v7.4, Stoelting Co., IL). Social preference was quantified as the interaction ratio, calculated by dividing the time spent in the chamber containing the novel conspecific by the total time spent in both outer chambers.

#### Open field test

Mouse was placed in an acrylic open-field arena (43 × 43 × 43 cm) virtually divided into a peripheral zone and a central zone (15 × 15 cm) for 15 minutes. Total time spent in the central zone (26), recorded using ANY-maze 7.4 video tracking software, was used as an index of anxiety-like behavior, with reduced center time indicating increased anxiety. Total distance traveled within the arena was measured as an indicator of locomotor activity.

#### Elevated plus maze test

This consisted of a plus-shaped acrylic apparatus elevated 94 cm above the floor, with two open arms (30 × 5 cm) and two closed arms (30 × 5 × 16 cm) connected by a central platform (5 × 5 cm). Mouse was placed in the center of the apparatus facing an open arm, and behavior was recorded for 5 minutes (26). Total arm entries and time spent in and exploring the open arms were tracked using ANY-maze 7.4 video tracking software. Decreased time spent in the open arms was interpreted as heightened anxiety-like behavior.

#### Splash test

The dorsal coat of mice was sprayed three times with a 10% (v/v) sucrose solution and immediately placed in an empty 4-liter PYREX glass cylinder (15 cm diameter). Behavior was recorded for 5 minutes using a CCTV camera (Model -WV-CP294), and total grooming duration was manually scored by a blinded experimenter using Kinoscope open-source software (27).

#### Pup retrieval test

Testing was conducted in a quiet behavior room during the morning light cycle. Prior to testing, dams in their home cages were habituated to the testing environment for 5 minutes. Before the start of the test, the total number of live pups per dam, including any litter mortalities recorded from the day of birth (P0), was documented. Following habituation, dams were temporarily removed from the home cage, during which pups were placed individually at designated positions approximately 15 cm away from the nest site. The dam was then returned to the cage, and behavior was recorded for 10 minutes (28). Latency to retrieve first pup, and all pups to the nest was manually scored by a blinded experimenter. Pups not retrieved within the 10-minute observation period were assigned a maximum latency of 600 seconds.

#### Y-maze barrier task

This task components are broken down into 1) Habituation (2 days, 15 minutes each) - Prior to habituation, all mice were weighed to maintain body weight at approximately 90% of free-feeding weight. To establish familiarity with the reward, a small scoop of 20 mg sucrose-based food pellets (Bio-Serv) was placed in each home cage. 2) Forced-Choice Training (3 days, 10 trials/day, 1 minute/trial) - After habituation, Mice were randomly assigned to left or right arms as high reward (HR; 4 pellets) or low reward (LR; 2 pellets), counterbalanced across groups. In each forced-choice trial, one arm was blocked, alternating between HR and LR. Each session included five HR and five LR trials. Mice completed five HR and five LR forced-choice trials per session. 3) Free-Choice Training (3–5 days, 10 trials/day, 2 minutes/trial) - Each free-choice session began with one HR and one LR forced-choice trial to reinforce arm-reward associations. Mice then completed 10 free-choice trials; arm selection was recorded. Upon entering an arm, a divider contained the mouse until all pellets were consumed. Mice selecting the HR arm in ≥70% of trials (≥7/10) advanced to barrier testing. Omissions were noted when no arm was selected, or pellets were not fully consumed. Training continued until all mice met criterion. 4) Barrier Testing - On test days, mice completed 3–5 forced-choice trials for each arm. A 10 cm inclined barrier was placed midway down the HR arm, followed by 10 free-choice trials. Latency and arm choice were recorded. Barrier testing was repeated on days 2 and 3 with 15 cm and 20 cm barriers, respectively (29).

#### RNA isolation

At P35, behavior-naive adolescent mice were weighed prior to tissue collection. A minimum of four mice per group and sex from mixed litters were used (Vehicle: Females n = 5, Males n = 6; prenatal THC: Females n = 5, Males n = 8; prenatal Stress: Females n = 5, Males n = 6; prenatal THC/Stress: Females n = 7, Males n = 4). Tissue punches of PFC and NAc were collected between 10:00 AM and 12:00 PM as previously described (33) and stored at −80 °C until ready for RNA extraction. Total RNA was isolated via Trizol (Invitrogen) homogenization and chloroform phase separation, followed by purification using the RNeasy Mini Kit (#74104, Qiagen) with a DNase step (#79254, Qiagen). RNA concentration and purity were assessed using a Nanodrop spectrophotometer (ThermoScientific).

#### Nanostring

RNA quality was assessed using the RNA 6000 Nano assay on an Agilent 2100 Bioanalyzer (Agilent Technologies, Santa Clara, CA) (36), and RNA integrity numbers (RIN) were generated using the 2100 Expert Software. Only samples with RIN > 8 were used for downstream transcriptomic analysis. Gene expression was quantified using the Nanostring nCounter Analysis System (Bruker, Seattle, WA) (30) with a custom-designed codeset targeting 337 genes. For each sample, 100 ng of total RNA was hybridized to reporter and capture probes according to the manufacturer’s protocol. Samples were processed in batches of 12 using the Nanostring Prep Station, and digital counts were acquired on the nCounter Digital Analyzer. Raw expression data were normalized and analyzed using nSolver Analysis Software (Nanostring Technologies).

#### Prenatal THC exposure

THC was prepared by dissolving 100 mg/ml THC in ethanol, followed by emulsification in 2% Tween-80, sonication, and dilution in physiological saline (24). All dams received daily s.c. injections of THC (2 mg/kg, 2 ml/kg) or vehicle from GD3 to birth.

#### Perinatal fentanyl exposure (PFE)

Detailed experimental description of the PFE paradigm has been discussed elsewhere (25, 31). In brief, pregnant dams were administered 10ug/ml fentanyl citrate in drinking water from GD3 through birth till weaning at postnatal day 21.Following weaning, juvenile pups were grouped housed by sex until ready for transcriptome profiling at adolescent age (postnatal day 35).

### Statistics

All statistical analyses were performed using GraphPad Prism (version 10.4.1). Behavioral data from both dams and adolescent offspring were analyzed using two-way analysis of variance (ANOVA) with MWDS/no MWDS and vehicle/THC exposure as between-subject factors. For adolescent behavioral assessments, males and females were analyzed separately to examine sex-specific effects. Where significant interactions or main effects were detected, Dunnett’s post hoc tests were used to compare each experimental group to the vehicle-treated control group. For all analyses, statistical significance was set at p < 0.05. Data are reported as mean ± SEM, with exact F- and p-values presented in the Results. Sample sizes are indicated in figure legends, and assumptions of normality and homogeneity of variance were confirmed prior to analysis. For gene expression analysis, normalized RLF files were analyzed using nSolver and GraphPad Prism (v10). Gene annotations done using Genecard resource, and ontology analysis was performed using Metascape (32) and ChEA3 (33).

## RESULTS

### THC and psychosocial stress during pregnancy disrupt maternal behavior and early postnatal outcomes

To model clinically relevant patterns of concurrent psychosocial stress and cannabis use during pregnancy, 8-week-old pregnant C57BL/6 mice were exposed to MWDS for 10 consecutive days, followed daily with subcutaneous injections THC (2 mg/kg) or vehicle injections until birth (Fig. 1A). Between gestational days 13-15, dams were evaluated for anxiety-related behaviors including social interaction (SI) test, open field test (OFT), and elevated plus maze (EPM). In the SI test (Fig. 1B), dams exposed to THC, MWDS, or their combination spent significantly less time interacting with conspecific compared to vehicle exposed dams, indicative of increased social withdrawal. In the OFT (Fig. 1C), all exposure groups showed significant reduction in time spent in the center zone compared to vehicle exposed dams, consistent with heightened anxiety-like behavior. Locomotion was not significantly altered in the OFT.(Fig. 1D) and during the SI test (Fig. S1). In the EPM (Fig. 1E), dams exposed to MWDS, with or without THC, spent significantly less time in the open arms. Notably, THC exposure alone did not alter EPM behavior. Postnatal assessments on day 5 reveal that both THC and stress exposures significantly increased litter mortality (Fig. 1F). Litter size was significantly reduced in all exposed groups relative to the vehicle group (Fig. 1G). In the pup retrieval test (Fig. H, I), we observed prolonged latencies to retrieve pups in all exposure groups compared to vehicle group. Specifically, all exposed dams exhibited delayed first pup retrieval (Fig. H), while only dams exposed to MWDS, with or without THC showed significant delays in full litter retrieval (Fig. I). These findings suggest stress and THC exposures impair early maternal responsiveness, with additive effects observed when both exposures occur concurrently.

### Prenatal stress, THC, and combined exposures induce sex-specific deficits in adolescent anxiety-like and motivated behaviors

To examine long-term behavioral consequences of prenatal exposures, adolescent offspring (P35–P45) were assessed for anxiety-like, and effortful motivated behaviors. In the SI test (Fig. 2A,A’), both male and female offspring exposed to prenatal stress or combined THC/stress displayed reduced social interaction, while prenatal THC exposure alone had no significant effect. In the OFT (Fig. 2B,B’), all male exposure groups showed reduced center zone exploration, indicative of increased anxiety-like behaviors. In females, this anxiogenic effect was limited to the prenatal stress and THC/stress groups, while THC alone had no effect. Importantly, total locomotion remained comparable across groups and sexes (Fig. 2C,C’). On the EPM (Fig. 2D,D’), both sexes exhibited reduced open arm exploration following combined prenatal THC/stress exposure. Males were particularly sensitive to prenatal stress exposures alone, while females showed heightened anxiety only following THC or combined exposures. The splash test (Fig. 2E,E’) revealed reduced grooming behavior, a proxy for self-care motivation in both sexes following prenatal THC or THC/stress exposures. Prenatal stress exposure alone also impaired grooming in females but not males. In the Y-maze barrier task, a measure of effort-based reward motivation, prenatal stress and THC exposures reduced HR arm selection under increasing effort demands. In the 10 cm barrier (Fig. 2F,F’), both sexes exhibited reduced HR-arm preference following combined THC/stress exposure, while prenatal stress only males and THC only females also showed decreased effortful motivation for sucrose reward. In the 15 cm barrier (Fig. 2G, G’), HR-arm selection reduced across groups, especially in males.

**Figure 2.**
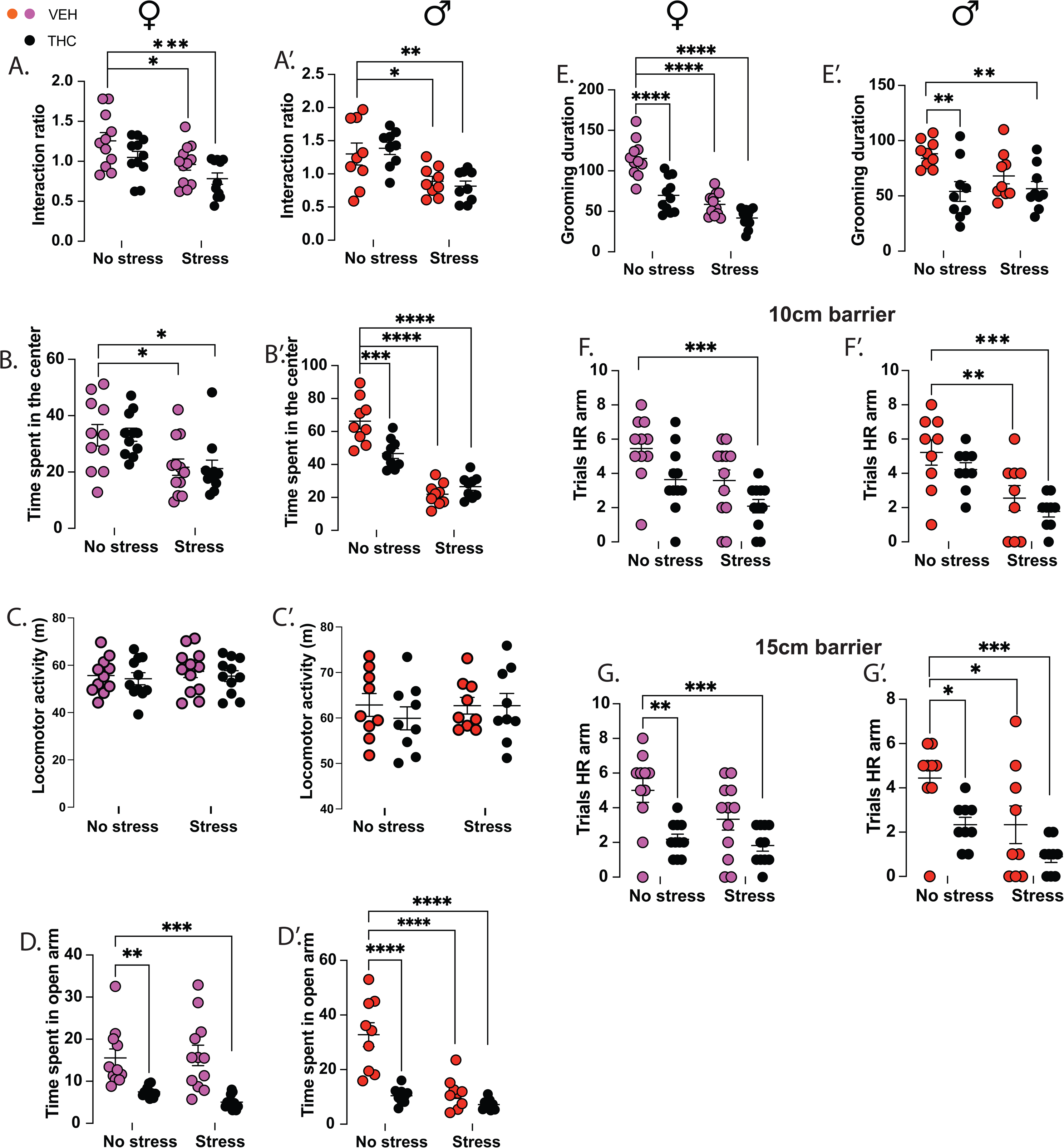
Concomitant exposures to prenatal THC and stress heightens anxiety- and motivation-like behaviors at adolescence. Social interaction test in (A) females and (A’) males, Open field test in (B) females and (B’) males, Elevated plus maze test in (C) females and (C’) males, Splash test in (D) females and (D’) males, Ymaze barrier (10cm) task in (E) females and (E’) males, and Ymaze barrier (15cm) task in (F) females and (F’) males. In the SI test (Fig. 2A, A’), two-way ANOVA revealed no interaction between prenatal THC- and stress-exposure in either sex (females: F_1,41_ = 0.03, p = 0.86; males: F_1,32_ = 0.54, p = 0.47). However, main effects of prenatal stress was observed in both females (F_1,41_ = 11.95, p = 0.0013) and males (F_1,32_ = 20.32, p < 0.0001), and a main effect of prenatal THC in females only (F_1,41_ = 5.607, p = 0.02; males: F_1,32_ = 0.002, p = 0.9). Post hoc analysis showed reduced interaction times in prenatal stress- (females: p = 0.03; males: p = 0.03) and THC/stress-exposed groups (females: p = 0.0006; males: p = 0.009), while prenatal THC exposure alone had no effect in either sex (females: p = 0.2; males: p = 0.8). In the OFT (Fig. 2B, B’), an interaction between prenatal stress- and THC-exposure was detected only in males (F_1,32_ = 4.46, p < 0.0006), but not females (F_1,41_ = 0.01, p = 0.9). Main effects of prenatal stress-exposure were present in both sexes (females: F_1,41_ = 14.65, p = 0.0004; males: F_1,32_ = 103, p < 0.0001), while a prenatal THC main effect was found only in males (F_1,32_ = 5.621, p = 0.02; females: F_1,41_ = 0.01, p = 0.9). Post hoc analyses revealed reduced center time in all male prenatal exposure groups (THC: p = 0.0004; stress: p < 0.0001; THC/stress: p < 0.0001), while in females, only the prenatal stress (p = 0.02) and THC/stress exposed (p = 0.02) groups showed significant reductions in time spent in center of arena. Prenatal THC-exposure did not alter center time in females (p > 0.9). Total locomotion in the OFT (Fig. 2C, C’) showed no interaction effects (females: F_1,41_ = 0.01, p = 0.89; males: F_1,32_ = 0.37, p = 0.54), nor main effects of prenatal stress- (females: F_1,41_ = 0.34, *p* = 0.56; males: F_1,32_ = 0.293, *p* = 0.592) or THC-exposure (females: F_1,41_ = 0.475, *p* = 0.49; males: F_1,32_ = 0.37, *p* = 0.545). Post hoc tests confirmed comparable total distance traveled across all groups. In the EPM (Fig. 2D, D’), an interaction between prenatal THC- and stress-exposures was found in males (F_1,32_ = 13.49, p = 0.0009), but not females (F_1,41_ = 0.8, p = 0.37). Main effects of prenatal THC were detected in both sexes (females: F_1,41_ = 32.13, p < 0.0001; males: F_1,32_ = 28.69, p < 0.0001), while a prenatal stress effect was present only in males (F_1,32_ = 24.78, p < 0.0001; females: F_1,41_ = 0.29, p = 0.594). Post hoc comparisons showed reduced open arm time in prenatal THC- (females: p = 0.0051; males: p < 0.0001) and THC/stress-exposed mice (females: p = 0.0003; males: p = 0.0001) Additionally, prenatal stress-exposure alone decreased open arm exploration in males (p = 0.0001), but not females (p = 0.988). To investigate motivated grooming behaviors, mice were tested in the splash test (Fig. 2E, E’). An interaction effect of THC × stress exposure was detected in females (F_1,41_ = 7.255, p = 0.01), but not in males (F_1,32_ = 2.673, p = 0.11). Main effects of prenatal THC-exposure were present in both sexes (females: F_1,41_ = 34.01, p < 0.0001; males: F_1,32_ = 10.9, p = 0.002), while prenatal stress-exposure had a main effect only in females (F_1,41_ = 62.61, p < 0.0001). Post hoc analyses showed reduced grooming time in prenatal THC-exposed (females: p < 0.0001; males: p = 0.004 and THC/stress-exposed mice (females: p < 0.0001; males: p = 0.0079). Stress alone decreased grooming time in females (p < 0.0001), but not males (p = 0.12). Effortful motivation for sucrose reward was evaluated using the Y-maze barrier task and we found in the 10 cm barrier phase (Fig. 2F, F’), no interaction of THC and stress in both sexes (females: F_1,41_ = 0.08, p = 0.76; males: F_1,32_ = 0.03, p = 0.849), but main effects of prenatal stress-exposure were found in both sexes (females: F_1,41_ = 9.664, p = 0.003; males: F_1,32_ = 19.41, p = 0.0001), and a prenatal THC main effect was present in females (F_1,41_ = 9.07, p = 0.004). HR-arm selection was reduced in prenatal THC/stress-exposed mice of both sexes (females: p < 0.0003; males: p = 0.0006), and in stress-only males (p = 0.0075) compared to vehicle-exposed mates. With the 15 cm barrier (Fig. 2G, G’), no interaction of THC x stress-exposures was observed (females: F_1,41_ = 1.577, p = 0.21; males: F_1,32_ = 0.35, p = 0.56). However, a main effect of stress was detected in males (F_1,32_ = 9.99, p = 0.003), but not females (F_1,41_ = 3.83, p = 0.057). Similarly, we observed main effect of THC in both sexes (females: F_1,41_ = 17.45, p = 0.0002; males: F_1,32_ = 9.99, p = 0.003). Post hoc analysis showed reduced HR-arm selection in THC- (females: p = 0.001 ; males: p = 0.03), THC/stress- (females: p = 0.0003; males: p = 0.0003), and stress-exposed males (males: p = 0.03 ; females: p = 0.06). By the 20 cm barrier phase, all groups failed to maintain HR-arm preference (data not shown), indicating a ceiling effect in task difficulty. Vehicle/no stress groups: females, n = 11, males, n = 9; THC/no stress groups: females, n = 11, males, n = 9; Vehicle/stress groups: females, n = 12, males, n = 9; THC/stress groups: females, n = 11, males, n = 9. *\*p =* 0.05, *\*\*p* = 0.01, *\*\*\*p* = 0.001, *\*\*\*\*p* < 0.0001.

### Prenatal THC and stress exposure drive brain region and sex specific transcriptional reprogramming in adolescent offspring

To uncover molecular correlates of behavioral changes, we conducted gene expression profiling using the Nanostring single molecular detection (nCounter) assay on PFC and NAc punches from adolescent male and female mice. The assay targeted 337 curated genes (Suppl. data 1) involved in synaptic function, neurodevelopment, mitochondrial metabolism, stress response, and immune signaling, previously found to be implicated perinatal fentanyl exposure (31). Principal component analysis (PCA) was first applied to reduce dimensionality and assess the contribution of sample characteristics to transcriptome variability and indicated that brain region was the primary driver of transcriptomic variance (Fig. 3A), with no distinct clustering by treatment or sex indicating minimal influence on sample variance. Treatment-driven effects were thus segregated by brain region. Broadly, prenatal stress induced upregulation of transcripts in the PFC and downregulation in the NAc (Fig. 3B). Volcano plots confirmed that differentially expressed transcripts (DETs) were primarily upregulated in the PFC with fewer downregulated DETs across all treatment groups (Fig. 3C, D). Venn diagrams show that seventeen and twelve DETs were shared across all three conditions in the PFC and NAc respectively (Fig. 3E, E’). Although sex did not emerge as a major contributor to global transcriptomic variance, we found sex-specific DETs not detected in combined-sex datasets (Fig. 3F, F’), underscoring distinct transcriptional signatures in males and females. Gene Ontology (GO) analysis showed that in the PFC (Fig. 3G), upregulated genes in the prenatal stress group were enriched for precursor metabolite and energy generation and transmembrane transporter complexes, indicating alterations in cellular metabolism and transport. In prenatal THC exposed mice, upregulated genes were enriched for synapse organization, excitatory synapse components, and semaphorin receptor activity, suggesting enhanced synaptic signaling and axon guidance processes. Combined THC/stress exposure led to upregulation of genes enriched for ribonucleoside triphosphate metabolism, respiratory chain complexes, and transmembrane transporter activity, reflecting increased bioenergetic demand. In contrast, downregulated genes in this group were enriched for cell–cell adhesion, indicating potential disruptions in structural connectivity. In the NAc (Fig. 3H), prenatal stress exposure resulted in upregulation of genes associated with metabolite and energy production and transcription factor binding, while downregulated genes were enriched for gliogenesis, cell adhesion, and integrin binding, suggesting suppression of glial and structural pathways. Prenatal THC exposure upregulated genes involved in metabolite generation, cation channel complexes, and ion channel activity, with downregulated genes enriched for gliogenesis, respiratory chain components, and integrin binding. In the Prenatal THC/stress group, upregulated genes were enriched for startle response, dendritic spine development, and glutamate receptor binding, while downregulated genes were associated with cellular respiration, plasma membrane signaling receptor complexes, and integrin binding, indicating altered synaptic excitability and cellular energetics.

**Figure 3.**
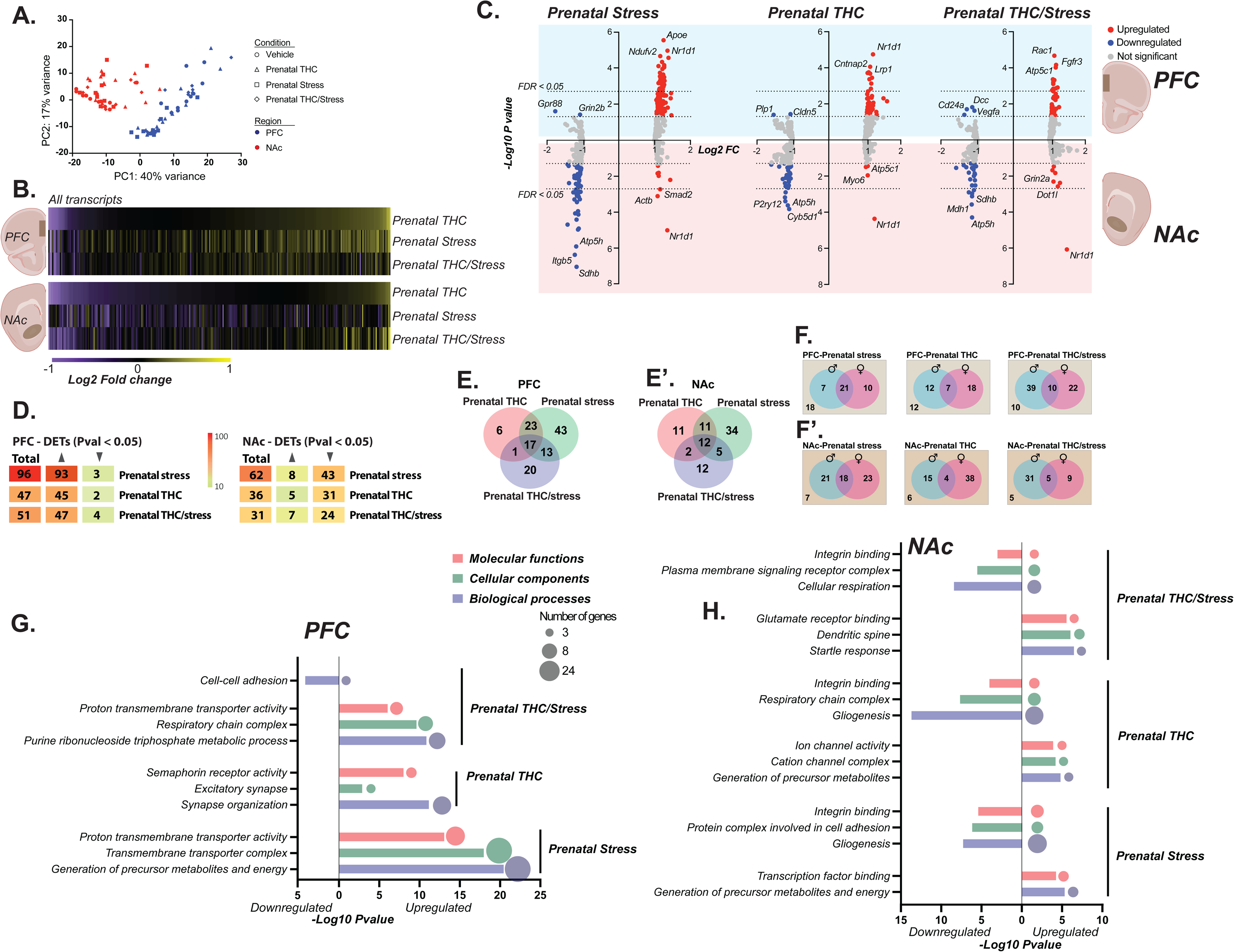
Transcriptome profiling in PFC and NAc of adolescent mice exposed to prenatal THC, stress and a combination of both. (A) Principal component analysis of PFC and NAc samples from all conditions. (B) heatmaps showing all gene transcripts (337) explored in the PFC and NAc of prenatal THC-, stress- and THC/stress-exposed adolescent mice. (C) Volcano plots showing DETs in the PFC (above) and NAc (below) across all exposed groups. Dotted lines on x-axis represent significance cut-off - *Pval* of 0.05 and FDR of 0.05. Red dots indicate upregulated transcripts, and blue dots indicate downregulated transcripts. (D) Shows total DETs and expression pattern in the PFC and NAc across the 3 exposed groups. Venn diagrams showing unique and shared DETs when sexes are combined in the (E) PFC and (E’) NAc and separated in the (F) PFC and (F’) NAc. Top gene ontology analysis in the (G) PFC and (H) NAc. Circle sizes indicate number of DETs present in enriched terms. Upregulated over-represented terms are shown to the right of vertical lines starting at zero, and downregulated over-represented terms on the left of vertical line. Bar graphs show biological processes (blue), cellular component (green) and molecular functions (red). DETs, differentially expressed transcripts.

To explore upstream regulatory mechanisms, we identified all significantly expressed transcription factors (TFs) across exposures and brain regions in a sex-wise manner using GeneCard annotations (Figs. 4, 5). In the PFC, the number of significant TFs in males vs. females were prenatal stress (1 vs. 8), prenatal THC (3 vs. 7), and prenatal THC/stress (4 vs. 6) (Fig. 4A). We compared our transcription factor list to ChEA3-predicted regulators (33), a tool developed based on RNA-seq and ChIP-seq datasets, to validate enrichment and support the functional relevance of TFs identified in our dataset. This revealed that in the PFC, upregulated DETs in prenatal stress-exposed males were enriched for regulation of secretion, with *Olig1* as the predicted TF (Fig. 4B). In females, enriched processes included anatomical structure maturation and regulation of neurogenesis, predicted to be regulated by *Myrf* and *Sox10* (Fig. 4C). *Sox10* was identified to be significantly upregulated in females in our dataset (Fig. 4A). In THC-exposed mice, DETs were enriched for cation transport in males with top TF *Hivep2* (Fig. 4D). Upregulated DETs in females were enriched in GABAergic neuron differentiation with top TF *Foxg1*,(Fig. 4E). Downregulated DETs in THC-exposed males were associated with cell-cell adhesion, with *Sox8* as the top TF (Fig. 4F). In THC/stress-exposed mice, upregulated genes were enriched in aerobic respiration with top predicted TF *Alx1* (Fig. 4G) while female upregulated DETs were enriched for epithelial cell proliferation with *Mterf2* as the predicted TF (Fig. 4H). Downregulated genes in THC/stress-exposed females were enriched for semaphorin-plexin signaling, with *Egr1* identified as the top regulator (Fig. 4I). Notably, *Egr3*, a related TF, was identified to be significantly downregulated in our dataset (Figs. 4A, I). In the NAc, the number of significant TFs in males vs. females were prenatal stress (5 vs. 4), prenatal THC (3 vs. 8), and prenatal THC/stress (3 vs. 2) (Fig. 5A). In stress-exposed females, upregulated DETs were enriched for memory formation with top predicted TF *Znf484* (Fig. 5B). In males, prenatal stress exposure downregulated genes enriched for myelin sheath development and predicted to be regulated by *Znf654* (Fig. 5C). In prenatal THC-exposed females, upregulated genes were associated with development and predicted to be regulated by *Lcor* (Fig. 5D). Downregulated genes in THC-exposed males were enriched in NAD metabolic process with *Znf654* as the top predicted TF (Fig. 5E) and in females, synaptic signaling process was top enrichment for downregulated DETs with *Nfiz* as predicted TF (Fig. 5F). In THC/stress-exposed mice, upregulated genes in males were enriched for cell-cell adhesion with *Ahdc1* as top predicted TF (Fig. 5G), and females show enrichment of DETs involved in response to growth factor with predicted TF *Myrf* (Fig. 5H). Downregulated genes in THC/stress-exposed females showed enrichment of cell-cell adhesion, with *Bcl6b* as the predicted TF (Fig. 5I).

**Figure 4.**
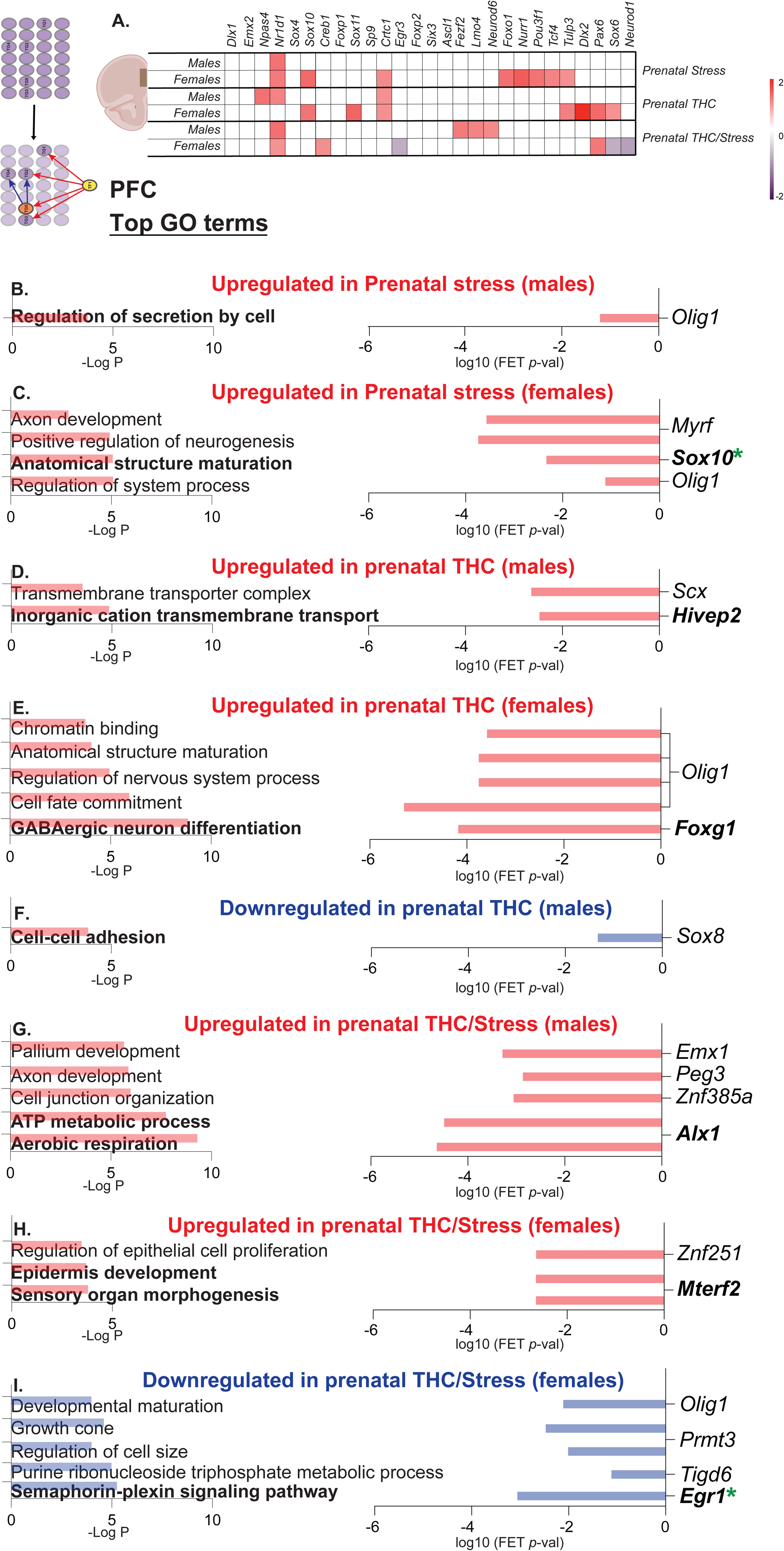
Transcription factor profiling in the PFC across exposed groups. (A) Table shows differentially expressed transcriptional regulators in the PFC by sex across exposed groups in our dataset. (B-I) shows top gene ontology enrichments of DETs on the left and predicted transcription factors in enriched terms on the right across groups. DETs, differentially expressed transcripts. FET, Fisher’s Exact Test.

**Figure 5.**
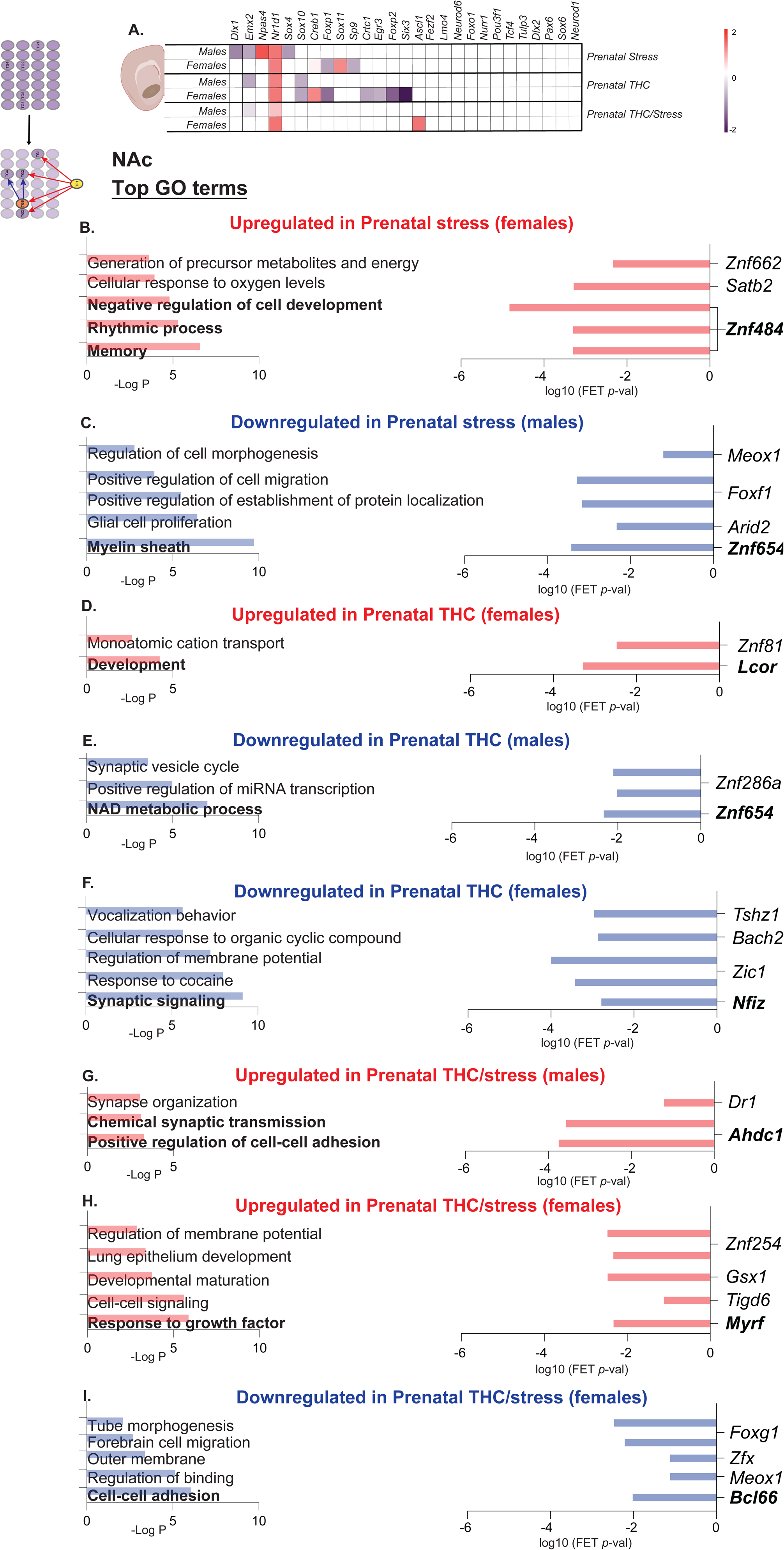
Transcription factor profiling in the NAc across exposed groups. (A) Table shows differentially expressed transcriptional regulators in the NAc by sex across exposed groups in our dataset. (B-I) shows top gene ontology enrichments of DETs on the left and predicted transcription factors in enriched terms on the right across groups. DETs, differentially expressed transcripts. FET, Fisher’s Exact Test.

### Cross-model transcriptomic convergence reveals common molecular signatures of early-life adversity

Developmental exposures to addictive substances or early life stress independently disrupts reward processing and increases vulnerability to negative affect. We build on our previous findings that low-dose of PFE induces persistent affective deficits and mesolimbic circuit disruption (25). We investigated whether shared behavioral outcomes across prenatal THC, stress and fentanyl exposures reflect convergence at the transcriptomic level. Using our curated list of 337 genes, we compared gene expression patterns in the PFC and NAc across exposures and sex (Fig. 6). In the PFC, THC- and stress-exposed females showed general transcript upregulation relative to PFE females and all male groups (Fig. 6A). In males, 5 DETs *Cspg5*, *Nr1d1*, *Ptger3*, *S1pr1* and *Syn3* were upregulated across all exposure groups (Fig. 6B, C). Four DETs overlapped between PFE and stress-exposed males including *Cd200*, *Mtfr1l*, *Mzt1*, and downregulated *Cldn5*, while *Shank1* was shared between PFE and THC-exposed males. Among females, four DETs, *Arhgef1*, *Atp5g2*, *Mtfr1l* and *Syn3* were shared across all exposures (Fig. 6D, E). Six DETs were shared between PFE and stress-exposed females, *Bin1*, *C1qc*, *Foxo1*, *Nr1d1*, *RhoA*, *Tmed7* and *Plxna3* was the only gene shared between PFE and THC in females. Notably, *Syn3* was upregulated across all groups and sexes in the PFC. Functional enrichment of shared DETs showed postsynaptic density and secretion regulation in males, and cell differentiation along with extrinsic component of membrane in females (Fig. 6F). In the NAc, a predominant downregulation of gene profile across all groups was observed (Fig. 6G). Four DETs *Cldn5*, *Hsp90*, *Lhx2* and *Nr1d1* were shared across all male exposures with *Nr1d1* consistently upregulated across exposures (Fig. 6H, I). Nine DETs were shared between PFE and prenatal stress-exposed males, with *Mzt1*, *P2ry12*, *Tprkb* and *Zfp748* showing opposing expression profile. No shared DETs were found between PFE and prenatal THC exposures. In females, 11 DETs were shared across all three groups, including six downregulated genes and four genes, *Atp5k*, *Dnm1l*, *Gpbp1*, *Ndufv2* upregulated in THC and stress groups. *Gria1* was uniquely upregulated by prenatal stress exposure but downregulated in prenatal THC and PFE. Ten DETs were shared between PFE and prenatal stress exposure with 5 DETs consistently downregulated in both exposures and 5 DETs upregulated only in prenatal stress exposure. Twelve DETs overlapped between PFE and THC with 11 DETs downregulated in both conditions, and 1 DET, *Atp5j2* upregulated only in prenatal THC exposure (Fig. 6J, K). Enrichment analysis of shared DETs reveal negative regulation of cell differentiation and regulation of protein localization process males, and ATP synthesis, gliogenesis and regulation of microglial migration in females (Fig. 6L).

**Figure 6.**
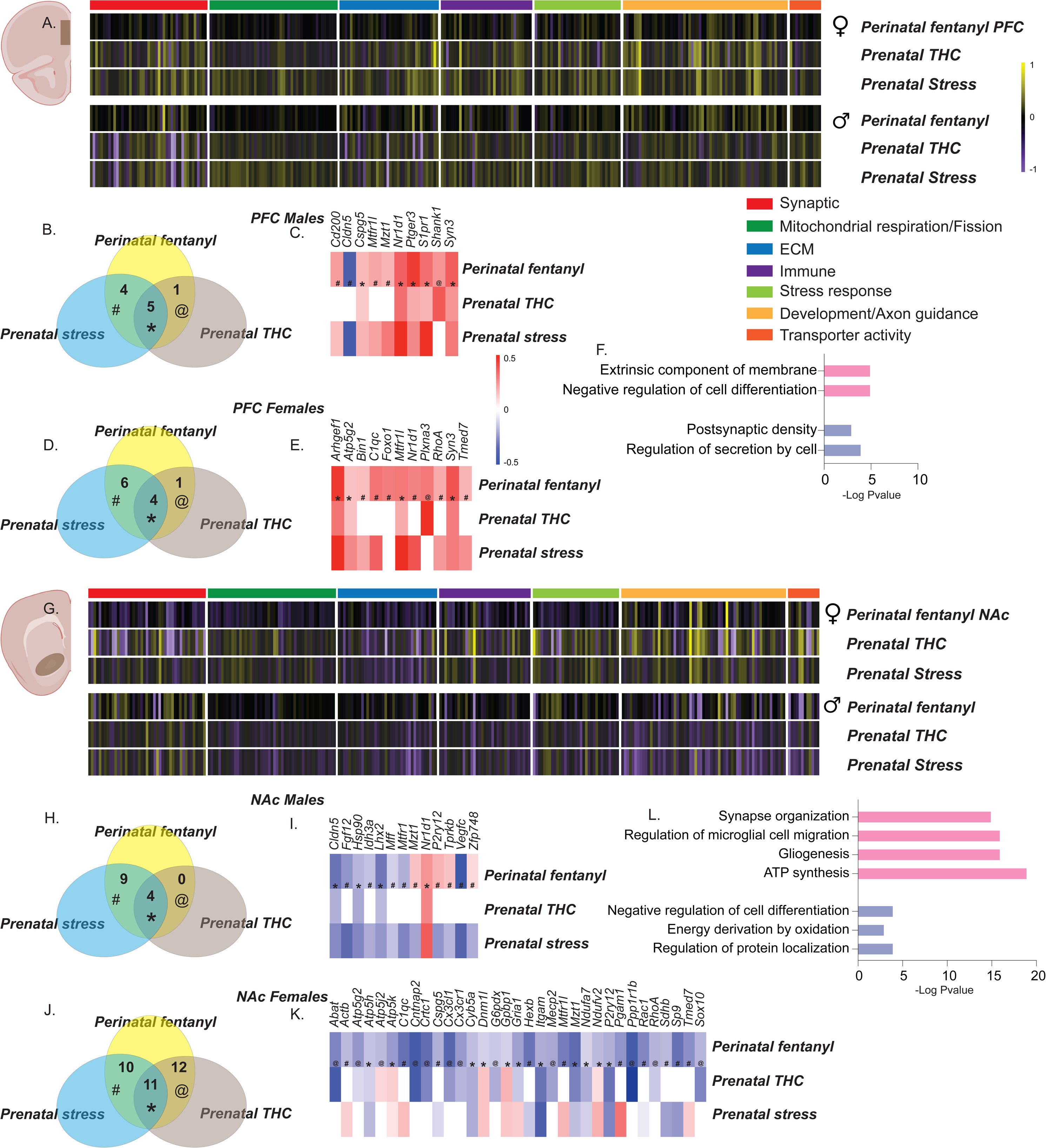
Transcriptomic convergence in the PFC and NAc of adolescent mice exposed to prenatal THC, stress and fentanyl. Heatmaps of all transcripts categorized as synaptic, mitochondrial respiration/fission, ECM, Immune, Stress response, development/axon guidance and transporter activity in the (A) PFC and (G) NAc, by sex of perinatal fentanyl-, prenatal THC- and stress-exposed adolescent mice. Venn diagrams show shared DETs in the PFC of (B) males and (D) females across conditions. Heatmaps show expression pattern of shared DETs in the PFC of (C) males and (E) females across conditions. (F) Gene ontology enrichment of shared DETs in the PFC of males (blue) and females (pink) adolescent mice. Venn diagrams show shared DETs in the NAc of (H) males and (J) females across conditions. Heatmaps show expression pattern of shared DETs in the NAc of (I) males and (K) females across conditions. (L) Gene ontology enrichment of shared DETs in the PFC of males (blue) and females (pink) adolescent mice. ECM, extracellular matrix. DETs, differentially expressed transcripts.

## DISCUSSION

The rising use of cannabis during pregnancy, often motivated by its perceived anxiolytic properties, raises serious concerns about long-term neurodevelopmental consequences in the offspring (34,35). Chronic psychosocial stress during gestation is a well-established disruptor of fetal brain development, acting either through direct alterations to neurodevelopmental trajectories (36,37) or by epigenetically priming neural circuits for heightened vulnerability to future stressors (38,39). Although findings on the effects of prenatal cannabis exposure are mixed (40,41), the critical role of the endocannabinoid (eCB) system in early brain development (42) suggests that cannabis use during gestation, particularly under conditions of chronic stress, may synergistically impair neurodevelopmental outcomes. Here, we established a novel rodent model combining prenatal THC exposure with chronic psychological stress to examine their additive effects on maternal behavior, adolescent behavioral outcomes, and transcriptomic signatures in the PFC and NAc. We employed the MWDS paradigm to model vicarious trauma, a form of psychosocial stress with translational relevance to human experiences of chronic threat or violence (20,43,44). Daily subcutaneous administration of 2 mg/kg THC from gestational day 3 to birth modeled moderate, clinically relevant exposure (45,46) while avoiding cannabinoid tetrad effects (47). Co-exposure exacerbated maternal caregiving deficits, as reflected by reduced litter size, increased pup mortality, and delayed pup retrieval. These findings challenge assumptions of THC’s anxiolytic benefit during pregnancy and suggest that combined THC and stress exposures disrupt maternal regulation.

Adolescent offspring exposed to combined prenatal THC and chronic stress exhibited robust impairments in anxiety-related, and motivated behaviors. Both sexes showed heightened anxiety-like phenotypes, consistent with studies indicating that prenatal THC sensitizes the HPA axis and increases emotional reactivity (48,49). Similarly, gestational stress alone disrupts glucocorticoid signaling and impairs brain development of emotional related circuitry (50,51), and our findings suggest that concurrent THC exposure exacerbates these effects. Social interaction deficits in both sexes align with evidence that prenatal cannabinoid exposure impairs sociability, likely via reduced endocannabinoid signaling and oxytocin function (52–54). These behavioral abnormalities are characteristic of early-life adversity and have been linked to persistent synaptic dysregulation in corticolimbic circuits (55–57). These findings support a “double-hit” model in which prenatal stress and cannabinoid exposure interact to potentiate long-term neurobehavioral dysfunction. Female mice exhibited more pronounced deficits in disrupted grooming behavior, consistent with human data showing a higher prevalence of depression among women (58,59). In contrast, male mice displayed more severe anxiety-like behaviors, a pattern that diverges from epidemiological trends. This discrepancy may reflect sex differences in symptom presentation or healthcare utilization, with males potentially underreporting or avoiding treatment for anxiety, thereby skewing population-level data (60).

To investigate molecular mechanisms underlying the observed behavioral deficits, we performed targeted transcriptomic profiling of 337 genes previously implicated in PFE (31), focusing on the PFC and NAc, regions central to executive function, emotion regulation, and reward processing (61,62). In males, combined THC/stress exposure upregulated *Mtfr1l* and *Atp5g2* in the PFC, suggesting disrupted mitochondrial fission and ATP synthesis. This aligns with reports that cannabinoids impair mitochondrial dynamics and neuronal energy balance (63,64). Enrichment of transcription factors *Olig1* and *Alx1*, involved in oligodendrocyte development and neurogenesis (65,66), may reflect heightened allostatic load impairing myelination during adolescence. In females, PFC upregulation of Fgf9, Pax6, and Jam2 suggests altered neurogenesis, cell fate, and BBB integrity (67–69). Downregulation of *Egr3*, a key regulator of synaptic plasticity and stress adaptation (70), alongside enrichment of semaphorin-plexin signaling, suggests impaired circuit refinement contributing to affective and motivational deficits (71). In male NAc, broad downregulation of genes related to cell adhesion and metabolism, including *Cldn5*, *Gria1*, *Eno3*, and *Mdh1*, was observed. Reduced *Cldn5* and *Gria1* implicate BBB dysfunction and impaired glutamatergic signaling, respectively (72,73), while *Eno3* and *Mdh1* downregulation signals compromised energy metabolism (74). Upregulated *Arhgef1* and enriched *Ahdc1* implicate disrupted cytoskeletal dynamics and neurodevelopmental vulnerability (75,76). In females, upregulation of *Ascl1* and *Fgf13*, genes regulating neurogenesis and sodium channel function (77,78) suggests altered excitability and differentiation of medium spiny neurons. Like males, *Eno3* and *Mdh1* downregulation in females highlights convergent mitochondrial dysfunction. Collectively, these findings underscore the importance of sex-specific investigations and support targeting mitochondrial and synaptic pathways for intervention. Public health messaging around prenatal cannabis use, particularly under stress, should consider these additive developmental risks.

Cross-model comparisons revealed convergent upregulation of *Syn3*, a key regulator of dopaminergic vesicle cycling (79,80), across perinatal exposures and sexes in the PFC, consistent with disrupted reward sensitivity and cognition in early-life adversity (81). In males, shared PFC upregulation of *Cspg5*, *Nr1d1*, *Ptger3*, and *S1pr1* suggests convergence on synaptic remodeling and circadian/metabolic pathways (82–84). Female PFC share upregulation of *Arhgef1*, *Atp5g2*, *Mtfr1l*, and *Tmed7*, reflecting structural and bioenergetic adaptations (85,86). *Plxna3*, a semaphorin receptor implicated in axonal guidance, was selectively upregulated in THC- and PFE-exposed females (87). In the NAc, male offspring shows shared downregulation of *Cldn5*, *Hsp90*, and *Lhx2*, with *Nr1d1* uniquely upregulated across all exposures. Reduced *Hsp90* and *Lhx2* suggest impaired glucocorticoid signaling and neurogenesis, respectively (88,89), consistent with reduced plasticity and reward motivation. In females, a broader set of 11 genes were shared across models, but with divergent regulation. Notably, *Atp5k*, *Dnm1l*, *Gpbp1*, and *Ndufv2* were downregulated in PFE but upregulated in THC and stress-exposed females, suggesting stimulus-specific metabolic reprogramming. *Dnm1l* and *Gpbp1* regulate mitochondrial fission and redox balance, respectively (90,91), implicating mitochondrial pathways as a sex-specific vulnerability node. GO enrichment confirmed sex-biased patterns: males showed enrichment in postsynaptic density and secretion regulation, while females exhibited enrichment in ATP synthesis, gliogenesis, and microglial migration (92), suggesting distinct compensatory strategies to prenatal insults. These findings emphasize that while both sexes exhibit shared molecular adaptations across diverse prenatal exposures, the directionality, cellular processes, and potential compensatory capacity differ markedly.

Our findings underscore the need for further translational research to explore the complex interplay between prenatal cannabis exposure and maternal stress given that many pregnant people report using cannabis for its perceived anxiolytic effects and stress relief.

## ACKNOWLEDGMENTS AND DISCLOSURES

This work was supported by the Matthew Osborne Foundation and the Kahlert Institute for Addiction Medicine [to JO]; Grant No. R01MH106500, R01DA054905, R33DA052101 [to MKL].

Conceptualization, JO, JC, KSM, MKL; Methodology, JO, MD, AK, MKL; Formal Analysis, JO, GK, DF; Investigation, JO, MD, MKL; Resources, JC, MKL; Writing-Original Draft, JO; Writing-Review & Editing, JO, JC, KSM, MKL; Visualization, JO; Supervision, JO, MKL; Funding Acquisition, JO, MKL.

The authors report no biomedical financial interests or potential conflicts of interest.

