## Supplementary Figure 1 for "Concurrent maternal stress and THC exposure during pregnancy alters adolescent behavioral outcomes and corticolimbic molecular programs"

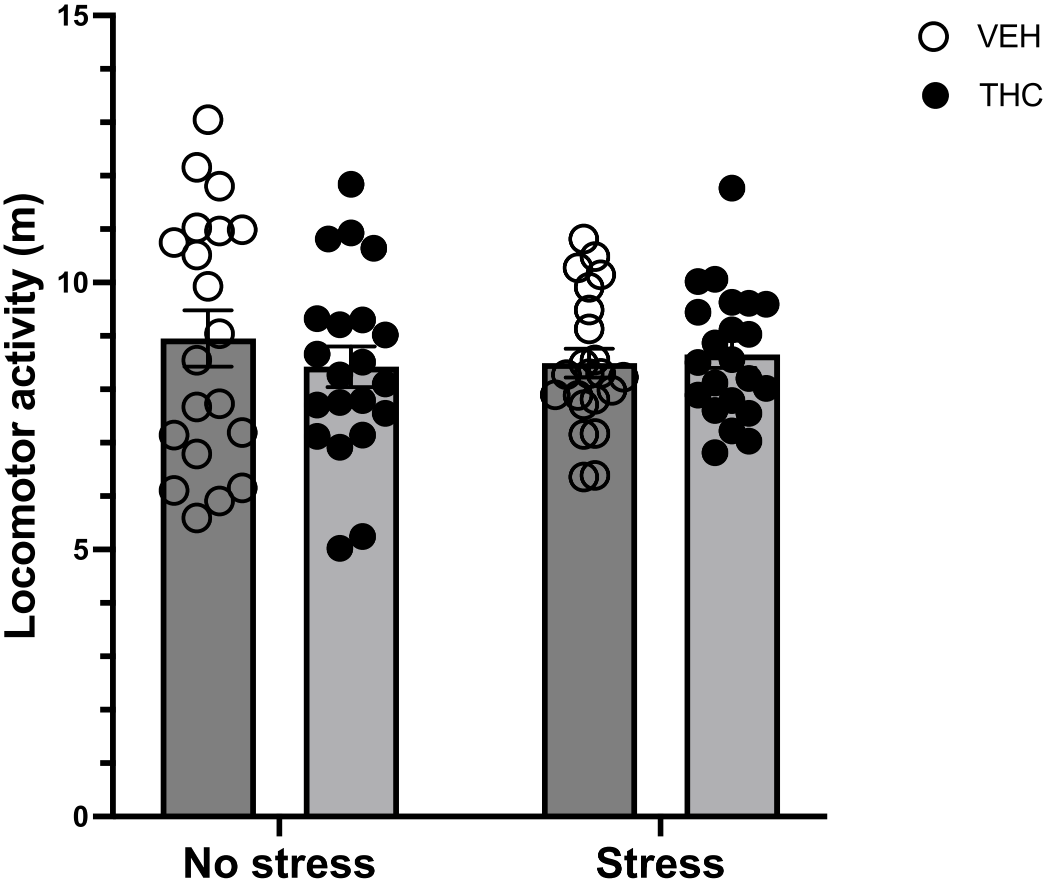


Fig. S1: Locomotor activity of dams in the three-chamber arena. All dams had comparable locomotor activity as the vehicle-exposed groups. There were no interaction effects ($F_{1,81}=0.91$; $p = 0.34$) or main effects of THC ($F_{1,81}=0.25$; $p = 0.62$) or stress exposure ($F_{1,81}=0.09$; $p = 0.75$) on locomotion.
